# CMPortal: a community-oriented, data-driven resource to inform protocol design for cardiac modelling from human pluripotent stem cells

**DOI:** 10.1101/2024.09.04.611313

**Authors:** Chris Siu Yeung Chow, Sophie Shen, Amy Hanna, Sumedha Negi, Clarissa L. L. Tan, Shaine Chenxin Bao, Chen Fang, James E. Hudson, Woo Jun Shim, Yuanzhao Cao, Nathan J. Palpant

## Abstract

Protocol design and benchmarking is central to optimising model development using human pluripotent stem cell derived cardiomyocytes (hPSC-CMs). By applying data mining to decades of research and hundreds of peer reviewed studies, we evaluate how protocol variables associate with common properties of cardiac functional and physiological maturation. This resource is publicly accessible through CMPortal, a community-oriented website that provides data-driven tools for researchers to navigate leverage decades of knowledge for benchmarking protocol designs and outcomes for their dedicated applications in developmental biology, disease modelling, and drug screening.

---

Human pluripotent stem cell-derived cardiomyocytes (hPSC-CMs) are a widely used model for cardiac research, enabling studies of development, physiology, disease mechanisms, and drug responses.^1^ However, maturing hPSC-CMs toward adult-like phenotypes remains an ongoing challenge for modeling adult cardiac biology.^2^ Despite development of maturation strategies involving specialized media, co-culture systems, and various types of matrix and biomaterials, optimizing protocol conditions suitable for modelling adult heart phenotypes remain challenging due to high variability in experimental design and outcomes.

Ewoldt et al.^3,4^ recently reviewed 300 study designs and demonstrate that while maturity outcomes have improved over time, no significant differences exist between maturation strategies. They highlight difficulties in parsing individual effects of protocol variables due to complex design variables and inconsistent reporting, drawing attention to the need for data-driven approaches and community-oriented benchmarking strategies to improve the accuracy and efficiency of hPSC-CM protocol design.

Here, we develop a data-driven platform that offers user-guided analyses of protocol features and their association with functional outcomes. Rather than assuming direct correlations exist between protocol factors and physiological maturation, we use an unsupervised approach that enables detection of bidirectionality and context-dependency allowing parameters to be independently assessed.

We first expanded the Ewoldt database^3^ to 322 studies, including 14 metabolic maturation studies and 8 publications to capture recent developments in the field. We also added protocol design features including cell line sex, ancestry, and transcriptomic markers of myofilament isoform switching. The final database has standardized nomenclature across 400 protocol features spanning five categories: experimental design variables (e.g., media composition factors, plating densities, matrix), analysis methods (e.g., imaging techniques, electrophysiology approaches), cell profiles (details about cell line including sex, ancestry, and source), study characteristics (e.g., publication year, journal), and measured outcomes (e.g., measured sarcomere length, contractile force, calcium handling, myofilament isoforms) (**Figure 1A**).

**Figure 1.**
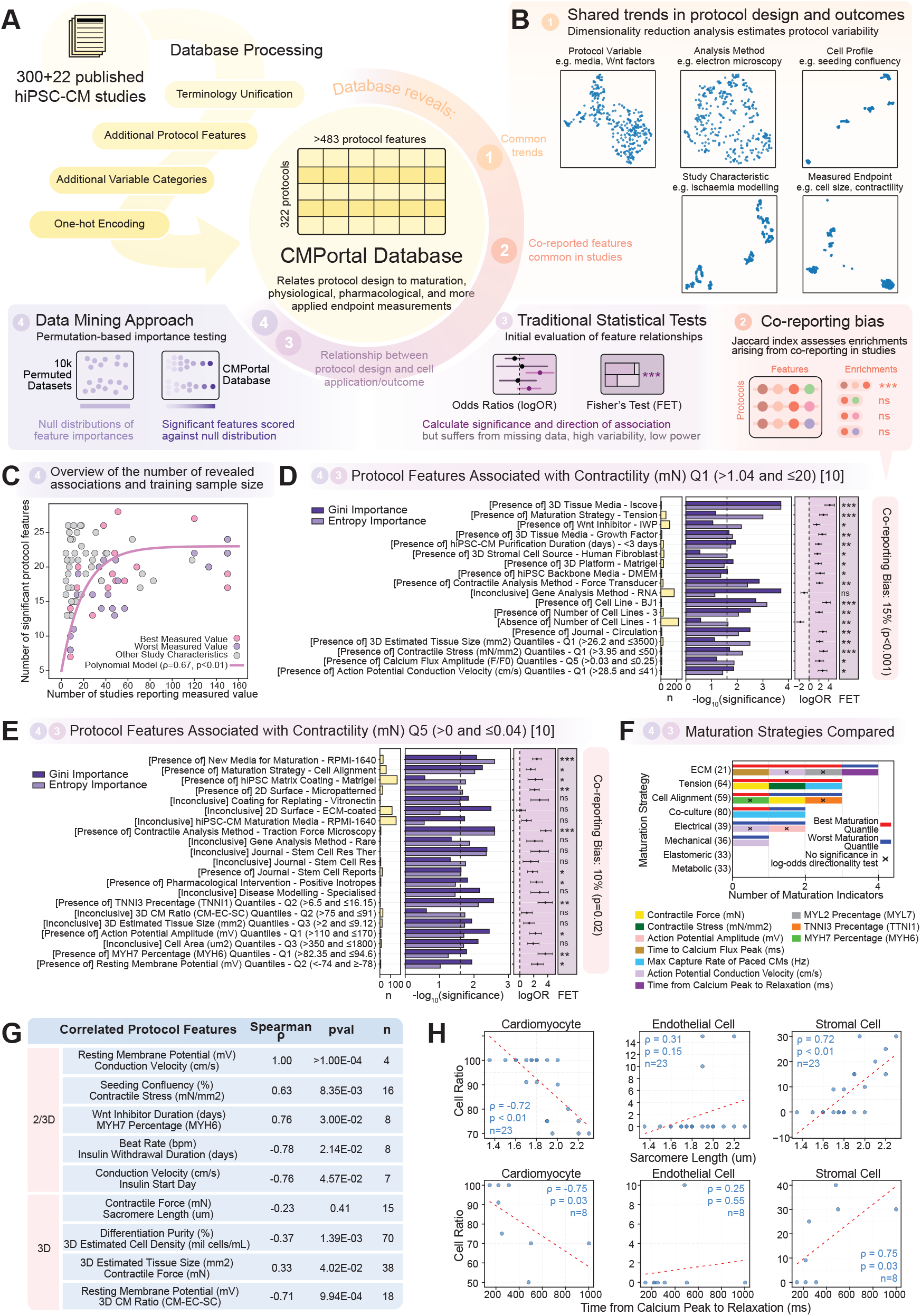
*CMPortal* database construction and hybrid data mining approach. **(A)** Schematic overview of the workflow for integrating and analysing 322 hPSC-CM studies to evaluate protocol designs and target parameters. **(B)** UMAP visualizations project the database by five feature categories, enabling assessment of variability in protocol components and identification of shared trends, with annotations shown in color. **(c)** Relationship between the number of significant enrichments and the number of training protocols. A polynomial model fitted to the best and worst quantiles of maturity indicators (pink and purple) suggests a plateau at 20–30 out of 413 protocol features significantly associated with maturity metrics. Grey indicates target parameters for intermediate maturity quantiles, cell profile, and study characteristics. **(D–E)** Bar plots showing protocol feature metadata for the highest (D) and lowest (E) contraction force quantiles, based on enrichments with p<0.025. Yellow bars indicate the number of studies reporting the relevant variables; purple bars represent the confidence in true feature enrichment. Log-odds ratio (logOR) and Fisher’s exact test (FET) estimate overall directionality of enrichment. Co-reporting bias is calculated as the percentage of enriched features co-published more often than expected by chance, using the mean Jaccard Index (JI) distribution for a matched number of features. **(F)** Stacked bar plot compares maturation strategies revealing significant context-dependent effects on maturation properties. **(G)** Table summarizing representative Spearman correlations among numerical protocol features, showing expected associations (full data in Table S4). **(H)** Spearman correlation plots of cardiac cell ratios in 3D cultures show opposing relationships with sarcomere length and calcium relaxation time as maturity indicators.

We applied a hybrid strategy combining conventional statistics and data mining to assess protocol features against 117 target parameters, such as 18 maturity indicators, cell profiles, and applications in disease and pharmacological modeling (**Tables S1 and S2**). Numerical measurements were binned into quantiles (e.g. Q1, Q2, etc.) to enable analysis of protocols ranked by their maturity outcomes. One-hot encoding was used to derive a data format suitable for machine learning, which also accounts for the significant missing data and assumes protocol variables, when not used or reported, are still potentially causal.

For data mining, we used random forest classifiers with 10,000 permuted feature sets to associate protocol designs with target parameters through Shannon entropy and Gini impurity, allowing the detection of statistically robust associations even in small and sparsely reported datasets (**Figure 1A** and **Table S3**). Dimensionality reduction via UMAP demonstrated high protocol heterogeneity across studies, consistent with reported findings^3^ (**Figure 1B**). Despite this heterogeneity, polynomial regression confirmed that the database sufficiently captures protocol features for target endpoints relating to best and worst performing quantiles in a data-limited setting (**Figure 1C**). Quality checks using Jaccard index analysis confirmed that data transformation preserved feature distinctiveness (**Figure S1**).

Data mining revealed alignment between protocol variables and expected biological impacts on cardiomyocyte maturation (**Figures 1D-E**). For example, Q1 contractility protocols were significantly associated with Wnt modulation, fibroblast inclusion, and matrix stiffness which are well established factors impacting sarcomere assembly and force generation.^5,6^ Importantly, we identified that identical maturation strategies could be linked to both best and worst maturation outcomes, demonstrating that this approach identifies context-dependent effects of protocol variables and not simply assume their generalized effects (**Figure 1F**). Specifically, the findings suggest that maturation parameters can be independently modulated by distinct sets of protocol variables, challenging the assumption that improving one cardiac maturity metric necessarily enhances all others.

We next used a spearman correlation analysis of all continuous variables in the database (n=31) to assess whether maturity measurements recapitulate expected physiological relationships (**Table S4**). Indeed, the data demonstrates that across hundreds of peer-reviewed studies, there are significant positive correlations between expected parameters including resting membrane potential and conduction velocity (ρ=1, p<0.01), as well as contractile force and 3D tissue size (ρ=0.33, p<0.05) (**Figure 1G**). Interestingly, the database also identifies protocol correlations that provide important considerations in co-culture design. Different cell type compositions revealed positive, null, and negative effects on maturity indicators including calcium kinetics and sarcomere length (**Figure 1H**).

We integrated these findings into a community-oriented web resource, *CMPortal* (https://palpantlab.com/cmportal). The Database Viewer enables searching of the 322 curated protocols based on five feature categories using techniques from simple keyword matching to multi-criteria searches, with options to toggle column visibility, export customized datasets, and evaluate relevant protocols of interest (**Figures 2A-C**). The Variable Search enables researchers to identify studies and protocols based on topic areas and/or sets of protocol variables which enables unsupervised discovery of relevant protocols based on desired and/or associated protocol features, even when the published protocols do not report the selected metrics (**Figure 2D**). The Protocol Benchmarking tool enables researchers to evaluate differentiation and maturation protocols against database standards. By uploading or manually selecting protocol features and the user’s own experimental measurements, the tool compares protocols through a radar chart that highlights maturity indicators across multiple categories and visually distinguishes between predicted, normal-range, and outlier experimental values (**Figures 2E-F**). Lastly, the Enrichment Browser allows researchers to identify protocol features significantly associated with specific outcomes which serves as the data foundation for the search and benchmarking features. (**Figure 2G**).

**Figure 2.**
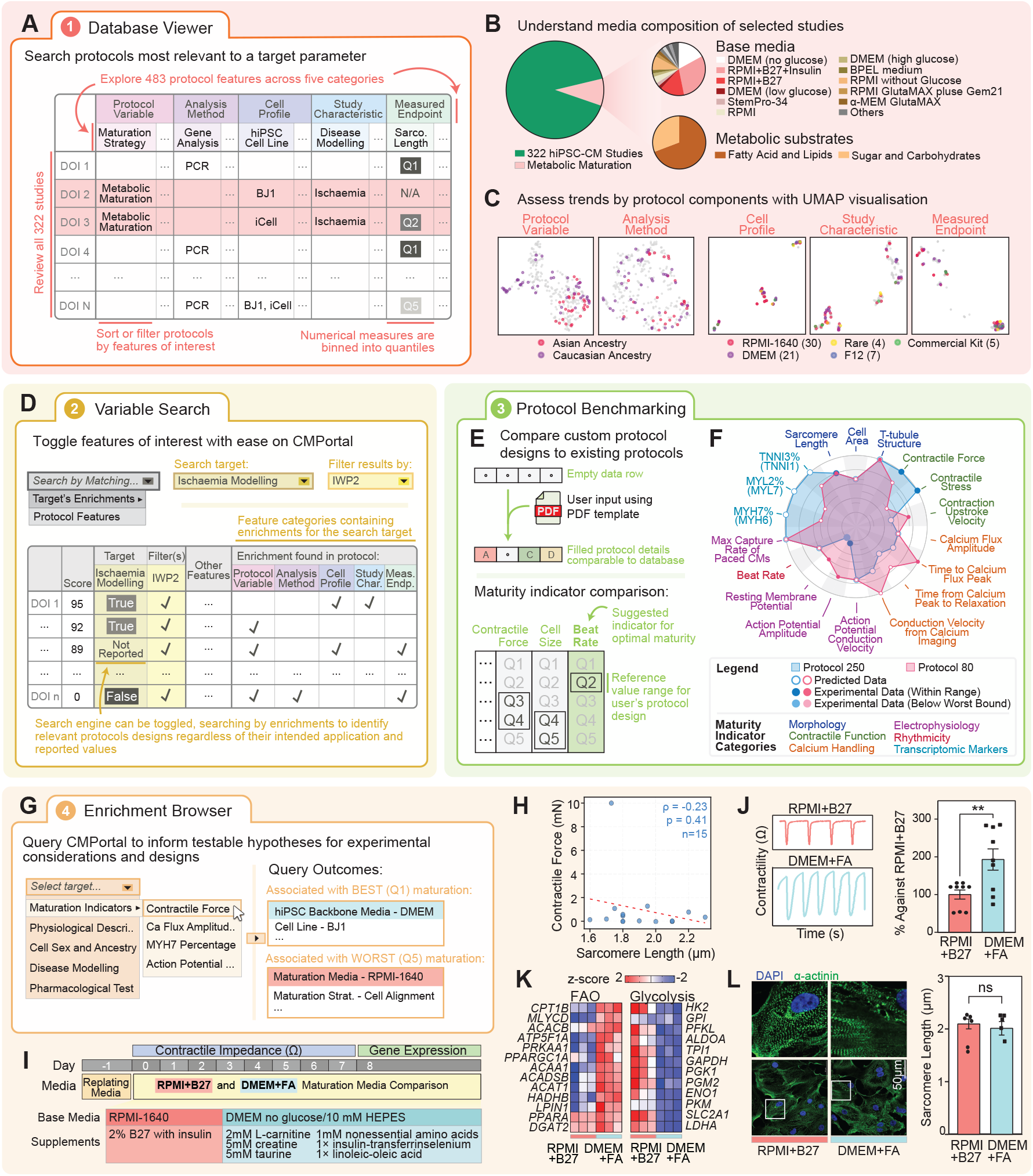
*CMPortal* website interface and experimental use cases. **(A)** Database Viewer allows users to explore 322 hPSC-CM protocols across 483 protocol features, and to install the filtered database. **(B)** Pie charts of the database show media and substrate usage of metabolic maturation protocols. **(c)** Example UMAPs show segregation of studies by the protocol feature categories. Cell line ancestry shows distinct trends of protocol variable and analysis method usage. Common media, like most features, show no clear trends. **(D)** Variable Search enables protocols retrieval by any desired and/or significantly enriched features by target parameters. **(E)** Protocol Benchmarking module enables users to upload and compare protocols against the database using experimental data and reference ranges estimated with the feature enrichments. **(F)** Radar plot generated by *CMPortal* for comparing maturation outcomes between Protocol 80 and 250 which are amongst the most cited studies in the database. The plot shows protocol 80 being associated to properties of higher transcriptional and contractile maturation, but lower electrophysiological and calcium handling maturation. Hallow datapoints are estimated with enrichments when no experimental data is available. **(G)** Enrichment Browser retrieves enrichments across the 117 target parameters. **(H)** Spearman correlation of pairwise, co-reported values of contractility and sarcomere length in 322 protocols. No significance was found. **(I)** *CMPortal*-informed experiment designed to compare the impact of media composition on contractility. **(J)** Representative traces and bar plot of impedance-based contractile measurements in hPSC-CM across time show higher peak amplitude for DMEM+FA and RPMI+B27 cells. **(K)** Bulk RNA-seq analysis of pathway-specific gene panels indicates metabolic reprogramming towards higher maturity in hPSC-CMs cultured in DMEM+FA compared to RPMI+B27. **(L)** Representative alpha-actinin staining (green), DAPI (blue) in fixed hPSC-CM confirms independence of contractility from sarcomere length. Bar charts values are mean ± standard error whiskers with student’s *t*-test significance (G, I). ^*^P<0.05, ^**^P<0.01,^***^P<0.001, ^****^P<0.0001, ns: non-significant. Study Char. : Study Characteristic; Meas. Endp.: Measured Endpoint; Sarco. Length: Sarcomere Length.

We aimed to validate an unexpected finding from the database showing that contractility and sarcomere length are not correlated maturation endpoints (**Figure 2H**), each having distinct protocol variable enrichments governing their maturation (**Table S2**). This finding suggests that the Frank-Starling effect, one of the most fundamental principles governing the relationship between heart pump function and load, develops through independent mechanisms. We investigated this by first identifying enriched protocol characteristics of Q1 and Q5 contractility, choosing backbone media compositions involving DMEM+fatty acids (FA) (Q1 contractility) and RPMI+B27 (Q5 contractility) (**Figure 2I**). DMEM+FA confers enhanced contractile maturity due to the high calcium concentration^7^ and fatty acids^5^ that mimic postnatal development. As shown in data mining, impedance-based measurements showed significantly higher peak contraction amplitude in DMEM+FA versus RPMI+B27 (**Figure 2J**), which was also supported by RNA-seq data showing higher metabolic activity from glycolysis to mature fatty acid oxidation in DMEM+FA cells (**Figure 2K**). Consistent with the database correlation, contractility changes occurred without significant variation in sarcomere length (**Figure 2L**). This supports our observation that fundamental features of cardiac maturation, including interdependent mechanisms controlling heart function like the Frank-Starling relationship, are not necessarily mechanistically linked in development. *CMPortal* allows for unsupervised discovery of these context-dependent protocol variables, enabling users to independently optimize protocols for specific functions.

*CMPortal* represents the first quantitative approach to overcome significant analytical difficulties of unreported, sparse protocol data, successfully associating context-dependent protocol variables with cell modelling outcomes. These findings challenge a fundamental but rarely challenged assumption that hPSC-CM maturity advances uniformly across all metrics. Instead, the results support a modularised paradigm to develop protocols through selected metrics based on experimental goals, aligning with recent literature demonstrating that attributes of hPSC-CM maturation can be compartmentalized.^8^ We translate these findings into user-friendly tools provided on a public website for researchers to evaluate, design, and benchmark protocols. *CMPortal* provides a streamlined approach to literature review, inform experimental design, and facilitate protocol benchmarking for understanding cardiac biology, diseases, and drug responses as a community-oriented resource for hPSC-CM research.

## Acknowledgements

This work has been supported by grant funding from the NHMRC (MRFCDDM000033 and 2007625 to NP), the Ian Potter Foundation (31111380 to NP), and the National Heart Foundation of Australia (106721 to NP). J.E.H. acknowledges the support of a Snow Medical Research Foundation Fellowship, Grant No. SMRF2019–060. Microscopy was performed at the Institute for Molecular Bioscience Microscopy Facility which was established with the support of the Australian Cancer Research Foundation (ACRF). We thank Meredith Redd for her assistance with project design.

## References

1. Murry, C. E. & Keller, G. Differentiation of Embryonic Stem Cells to Clinically Relevant Populations: Lessons from Embryonic Development. Cell 132, 661–680 (2008).

2. Yang, X., Pabon, L. & Murry, C. E. Engineering Adolescence: Maturation of Human Pluripotent Stem Cell– Derived Cardiomyocytes. Circulation Research 114, 511–523 (2014).

3. Ewoldt, J. K. et al. Induced pluripotent stem cell-derived cardiomyocyte in vitro models: benchmarking progress and ongoing challenges. Nat Methods 22, 24–40 (2025).

4. Ewoldt, J. et al. Induced pluripotent stem cell-derived cardiomyocyte in vitro models: tissue fabrication protocols, assessment methods, and quantitative maturation metrics for benchmarking progress. 192537 bytes Dryad 10.5061/DRYAD.KSN02V7BH (2024).

5. Lyra-Leite, D. M. et al. A review of protocols for human iPSC culture, cardiac differentiation, subtype-specification, maturation, and direct reprogramming. STAR Protocols 3, 101560 (2022).

6. Lee, S. et al. Contractile force generation by 3D hiPSC-derived cardiac tissues is enhanced by rapid establishment of cellular interconnection in matrix with muscle-mimicking stiffness. Biomaterials 131, 111–120 (2017).

7. Tiburcy, M. et al. Defined Engineered Human Myocardium With Advanced Maturation for Applications in Heart Failure Modeling and Repair. Circulation 135, 1832–1847 (2017).

8. Fetterman, K. A. et al. Independent compartmentalization of functional, metabolic, and transcriptional maturation of hiPSC-derived cardiomyocytes. Cell Reports 43, (2024).

