## Supplementary material for "CMPortal: a community-oriented, data-driven resource to inform protocol design for cardiac modelling from human pluripotent stem cells": Methods File

### STAR★METHODS

#### KEY RESOURCES TABLE

| REAGENT or RESOURCE | SOURCE | IDENTIFIER |
|---|---|---|
| **Antibodies** | | |
| Phycoerythrin (PE)-conjugated sarcomeric α-actinin (SA) antibody | Miltenyi Biotec Australia Pty | Cat.#130-123-773 |
| PE-conjugated anti-human isotype (IgG) control | Miltenyi Biotec Australia Pty | Cat.#130-119-964 |
| Anti-α-actinin antibody (mouse monoclonal) | Sigma | Cat.#A7811 |
| Alexa Fluor 488-conjugated anti-mouse secondary antibody | ThermoFisher | Cat.#A-11001 |
| **Chemicals, Peptides, and Recombinant Proteins** | | |
| mTeSR™ Plus maintenance medium | STEMCELL Technologies | Cat. #100-0276 |
| Bovine serum albumin (BSA) | Sigma Aldrich | Cat. #A9418-50G |
| L-Ascorbic acid 2-phosphate sesquimagnesium salt hydrate (ascorbic acid, AA) | Sigma Aldrich | Cat. #A8960-5G |
| CHIR-99021 (CHIR) | STEMCELL Technologies | Cat. #72054 |
| XAV-939 | STEMCELL Technologies | Cat. #72674 |
| Vitronectin XF | STEMCELL Technologies | Cat.#07180 |
| Paraformaldehyde (4%) | Sigma Aldrich | Cat.#158127-5G |
| Saponin | Sigma Aldrich | Cat.#S7900 |
| Trypsin (2.5%) | ThermoFisher | Cat. #15090046 |
| Versene Solution | ThermoFisher Scientific | Cat. #15040066 |
| RPMI-1640 | ThermoFisher | Cat. #11875093 |
| DMEM no glucose | ThermoFisher | Cat.#11966025 |
| HEPES (10 mM) | Sigma Aldrich | Cat. #H0887-100ML |
| L-carnitine | Merck Life Science Pty Ltd | Cat. #C0283-5G |
| Creatine | Sigma Aldrich | Cat. #C0780-50G |
| Taurine | Sigma Aldrich | Cat. #T8691-25G |
| Nonessential amino acids (1 mM) | ThermoFisher | Cat. #11140050 |
| Insulin-transferrin-selenium (100X) | ThermoFisher | Cat.#41400045 |
| Linoleic-oleic acid (100X) | Merck Life Science | Cat.#L9655-5ML |
| Y-27362 | STEMCELL Technologies | Cat. #72308 |
| B27 supplement with insulin (2%) | ThermoFisher | Cat. #17504001 |
| Fetal bovine serum (FBS) | ThermoFisher | Cat. #10099141 |
| Tween-20 (0.1%) | Sigma Aldrich | Cat. #P7949-100ML |
| Donkey serum (10%) | Sigma Aldrich | Cat. #D9663-10ML |
| Triton X-100 (0.1%) | Sigma Aldrich | Cat. #T8787-100ML |

| | | |
|---|---|---|
| DAPI (1 µg/mL) | Sigma Aldrich | Cat. #D9542 |
| **Deposited Data** | | |
| Raw and processed RNA-seq data | This manuscript | GEO: GSE298573 |
| CMPortal dataset | This manuscript | https://palpantlab.com/cmportal |
| Code for database processing and enrichment analyses | This manuscript | Available upon request |
| Adult human heart data | Hahn et al., 2021 | DOI: 10.5281/zenodo.4114617 |
| hPSC-CM database | Ewoldt et al., 2024 | DOI: 10.5061/DRYAD.KSN02V7BH |
| **Experimental Models: Cell Lines** | | |
| WTC-11 hPSC line | Gladstone Institute of Cardiovascular Disease, UCSF | RRID: CVCL_Y803 |
| **Software and Algorithms** | | |
| Python | Python Software Foundation | v3.12 |
| Pandas | pandas.pydata.org | v1.3.5 |
| NumPy | numpy.org | v1.21.0 |
| SciPy Stats | scipy.org | v1.12 |
| Scikit-learn | scikit-learn.org | v1.5.2 |
| Flask | flask.palletsprojects.com | v2.0.1 |
| Gunicorn | gunicorn.org | v20.1.0 |
| Nginx | nginx.org | v1.27.4 |
| GraphPad Prism | GraphPad Software | v9.3.1 |
| STAR aligner | STAR | https://github.com/alexdobin/STAR |
| htseq-count | HTSeq-count software | https://htseq.readthedocs.io/ |
| EdgeR package | Bioconductor | https://bioconductor.org/packages/edgeR |
| ImageJ | NIH | https://imagej.net/ |
| FACSDiva software | BD Biosciences | https://www.bdbiosciences.com/en-au/products/software/instrument-software/bd-facsdiva-software |
| FlowJo software (Version: 10.6.2) | Tree Star | https://www.flowjo.com/solutions/flowjo |
| CardioExcyte Control96 (Version: 1.4.7.1) | Nanion Technologies GmbH | https://www.nanion.de/products/cardioexcyte-96/ |
| **Other** | | |
| 96-well culture plates | Nunc | Cat.#150318 |
| CardioExcyte NSP-96 plates | Nanion Technologies GmbH | Cat.#201001 |
| CardioExcyte 96 system | Nanion Technologies GmbH | https://www.nanion.de/products/cardioexcyte-96/ |
| 35 mm MatTek glass-bottom dish | MatTek | P35G-1.0-14-C |
| Zeiss LSM 880 confocal microscope | Zeiss | https://imb.uq.edu.au/microscopy-confocal-2 |
| FACS CANTO II system | Becton Dickinson | https://www.bdbiosciences.com/en-au/products/instruments/flow- |

| | | |
|---|---|---|
| | | cytometers/clinical-cell-analyzers/facscanto |
| LightSail | Amazon Web Services | https://aws.amazon.com/lightsail/ |
| RNeasy Mini Kit | QIAGEN | Cat. #74106 |

### RESOURCE AVAILABILITY

#### Lead contact

#### Materials availability

This study did not generate new unique reagents.

#### Data and code availability

Bulk RNA-seq has been deposited at the Gene Expression Omnibus with accession code GSE298573. The reviewer token is kzqvgquwhfatjcp.

The code underlying computational analyses will be made available upon request given their length. The code for the website is available at (https://github.com/phycochow/palpant-labsite).

### EXPERIMENTAL MODEL AND SUBJECT DETAILS

**Generation and maintenance of human cell lines**

All hPSC studies were carried out in accordance with consent from The University of Queensland's Institutional Human Research Ethics approval (HREC#: 2015001434). Cardiomyocytes (CMs) generated in this study were derived from the WTC-11 hPSC line (Gladstone Institute of Cardiovascular Disease, UCSF).[3,4] All WTC cells were maintained as previously described with slight adaptions.[5,6] Briefly, hPSC cells were maintained in mTeSR Plus medium with supplement (Stem Cell Technologies, Cat.#05825) at 37°C with 5% $CO_2$. Cells were cultured on Vitronectin XF (Stem Cell Technologies, Cat.#07180) coated plates (Nunc, Cat.#150318).

### METHOD DETAILS

**Database Curation**

The foundational database[1] was expanded to include 322 published studies with over 442 features. It incorporates 14 additional hPSC-CM studies using metabolites, and 8 studies published in the previous year, 2024, excluding all reviews, genetic models and regeneration studies. These recent papers were retrieved on PubMed using the terms "ipsc AND cardiomyocytes AND maturation AND disease modelling". Next, features were added and removed. First, omics, which listed the multi-omics tools that were applied, was removed due to its ambiguity on the scope and purpose of these analyses. Second, Sex and Ancestry of cell lines were manually added with hPSCReg.[2] Third, GEO accession number was included for protocols reporting expression data of myofilament isoform switching.

**Database Processing**

Inconsistent entries and typos were first manually standardised. Examples include Wnt inhibitor BMP4 annotated as BPM4. We retained unexpected protocol variables in the dataset, like the differentiation media, StemPro-34, used for hiPSC backbone media,[3] to investigate minority protocols and unexpected effects. Next, feature entries were categorised into additional annotations. For example, metabolic media supplements and drugs were grouped by their broader function and class, aiming to generalise findings and improve the interpretability of associations. Underreported entries with less than 3 occurrences were then classified as Rare/Specialised. Similarly, similar numerical values were binned into quantiles to improve robustness for the underreported maturation endpoints. This was done blinded prior to statistical tests and data mining enrichment.

**Database Transformation**

To utilise full protocol information, one-hot encoding was applied across protocol features, assuming the absence of reported variables are as important as their presence. Each entry category was transformed into a new feature with binary values. Second, the values of numerical features were categorised, aiming for the maximum amount of quantile bins with a similar (±2) protocols count. Finally, each quantile was stored as new features. This manual process was assisted through code (Python 3.12). An example of one-hot encoding is shown below:

<table>
<tr><th>Preprocessing State</th><th>Protocol Feature</th><th>Values</th></tr>
<tr><td>Ewoldt et al.</td><td>Maturity Indicator</td><td>0.25, 1, …, 20</td></tr>
<tr><td>This study</td><td>Maturity Indicator - Quantiles</td><td>Q1, Q2, …, Q3</td></tr>
<tr><td rowspan="3">One-hot Encoded</td><td>Maturity Indicator - Q1 (10 studies)</td><td>True, False, …, False</td></tr>
<tr><td>Maturity Indicator – Q2 (10 studies)</td><td>False, True, …, False</td></tr>
<tr><td>Maturity Indicator – Q3 (11 studies)</td><td>False, False, …, True</td></tr>
</table>

**Jaccard Index analysis**

A pair-wise distance matrix was constructed using Jaccard Index (JI) between the transformed protocol features. While most protocol features maintain substantial uniqueness (mean JI=0.0275) (Extended Data Fig. 1), certain feature subsets exhibited significant similarity patterns. Notable examples include perfect matches (JI=1.0) among various metabolic components such as Albumax, B27, vitamin B12, Biotin, etc. which showed consistent co-reporting across 3 independent protocols. These technical artefacts emerged due to repeated sampling during the blinded database processing. Recognising their impact on downstream enrichments, we developed a co-reporting bias test described below to inform the readership of the protocol diversity behind the enrichments.

**Direct Statistical Tests**

Log-odds ratios were computed to quantify effect directionality between protocol features and target parameters. Contingency tables (2x2) were constructed for each feature-target pair, 1 is added to all cells if there zero-cells are present. Significance was assessed using two-tailed Fisher's exact tests in scipy.stats (v1.12) due to its robustness with small feature sample sizes. Multiple testing correction, which compromises association discovery, was not applied to enable conservative comparison between conventional statistics and data mining strategy.

#### Model Fitting and Importance Metrics

The full dataset of 322 protocols was used for model fitting, aiming to predict best and worst outcomes of maturation indicators and physiological descriptions, cell characteristics, and applications for pharmacology responses and disease modelling. Sklearn[4] random forest package (v1.5.2) was used with default parameters except for n_estimators, which was increased from 100 to 150. All models were fitted to the full dataset to maximise detection of feature association with the target protocols. Two similar metrics were used to capture maximum associations noting the small dataset sizes: Shannon Entropy (information gain via the reduction of entropy by splitting the dataset with the protocol feature) and Gini Impurity (the likelihood that the target is miscategorised in a subset after splitting with the protocol feature). Both metrics were used despite minor differences in their behaviour because they captured biologically meaningful enrichments that were not found by the other. Together, an enrichment is more technically robust if found significant in both. Since the scores reflect the degree to which a feature separates outcomes (e.g. Q1 maturity vs other quantiles), our approach captures protocol designs with bidirectional, context-dependent effect. The association directionality is therefore estimated with the described log-odds test, which was also used for comparison.

#### Feature Selection Strategy

A permutation-based strategy was developed to find authentic associations. First, 10,000 permuted datasets were generated by randomly shuffling the values within each protocol feature across all 322 studies. Second, the permutated datasets were processed through the same pipeline as the original dataset. The resulting 10,000 sets of importance scores established null distributions for both Gini and entropy enrichments for each predicted feature. Lastly, p-value was measured as the probability that the permutated version of a feature resulted in higher importance score than the original.

#### Co-reporting Bias

The mean JI was calculated for each enrichment panel to quantify the experimental diversity of the features using the pair-wise JI distance matrix. The Measured Endpoint feature category (Table S4) was excluded from the enrichments for the calculation. Next, the null mean JI distribution was generated using 5000 random panels with a matching number of protocol features. We calculated p-value by assessing whether the random features are more co-reported than the enriched feature panel.

#### *CMPortal* Development

*CMPortal* was developed as an extension of the laboratory website for hiPSC-CM protocol analysis. The backend server system was built using Flask (v2.0.1), a Python (v3.12) web framework, with a Model-View-Controller (MVC) modular architecture that separates the data management, user interface, and control logic. Data processing was performed using pandas (v1.3.5) and NumPy (v1.21.0) libraries. The front end is developed using HTML5, CSS3, and JavaScript. It was deployed using Gunicorn (v20.1.0) and a Nginx (v1.27.4) reverse proxy using LightSail provided by Amazon Web Services.

**Protocol Search Algorithm**

To assist the retrieval of protocols, we decided on a straightforward algorithm to maintain interpretability. The query protocol is first added to the one-hot encoded dataset (Table S1). For each of the 322 protocols, we then calculate the number of shared protocol features, including mutual absences. Lastly, five additional columns are appended to indicate whether each feature category contains matching features that are present in both the query and reference protocols. *CMPortal* supports search by target parameters regardless of their intended application or measurements including 16 maturity indicators, cell sex and ancestry, and applications in disease and pharmacological modelling, using their significant enrichments.

**Reference Ranges for Maturation Indicators**

The benchmarking module evaluates user-submitted protocol design by comparing them against the enrichments of maturation indicators. We designed an algorithm similar to motif search[5] in bioinformatics. The algorithm rewards +2 points for direct target quantile association, +1 point for neighbouring quantile association, 0 for the next neighbours', and penalises -1 for the features associated to remaining quantiles. Users control the score calculations by selecting relevant feature categories. For example, by selecting Protocol Variable or Analysis Method, they can focus on experimental factors that are more likely causal of the measurements. The system automatically recommends the quantile with highest enrichment as the optimal reference range.

**Flow cytometry**

At the time of replating on day 15 of differentiation, a subset of cells (approximately $1 \times 10^6$) was set aside for flow cytometry analysis of cardiomyocyte purity of differentiated cell populations. Cells were fixed with 4% paraformaldehyde (Sigma Aldrich, Cat.#158127-5G), permeabilised in 0.75% saponin (Sigma Aldrich, Cat.#S7900), and labelled with Phycoerythrin (PE)-conjugated sarcomeric α-actinin (SA) antibody (Miltenyi Biotec Australia Pty, Cat.#130-123-773) or PE-conjugated anti-human isotype (IgG) control (Miltenyi Biotec Australia Pty, Cat.#130-119-964). Stained samples were analysed using a FACS CANTO II (Becton Dickinson) system with FACSDiva software (BD Biosciences). Data analysis was performed using FlowJo (v10.6.2), and cardiac populations were determined with population gating from corresponding isotype controls. Cell preparations with >80% sarcomeric α-actinin-positive CMs were used in all experiments.

**Replate of CMs and maturation induction**

Differentiated CMs were replated on day 15 of differentiation for *in vitro* hPSC-CMs maturation protocols. Cells were dissociated using 0.5% trypsin, stopped with Stop Buffer (1:1 FBS in RPMI-1640), filtered with a 100 µM strainer, replated at a density of $1.58 \times 10^5$ cells/cm$^2$ in Vitronectin XF-coated 96-well plates with replating medium (RPMI-1640, 2% B27 supplement plus insulin, 5% FBS, and 10 µM Y-27362) and cultured overnight. FBS and ROCK inhibitor-containing medium was replaced the following day (day 0) with RPMI+B27 medium (RPMI-1640 plus B27/insulin). From the replating day 0 to day 7 post-replate, different backbone media were used to maintain CMs including RPMI+B27 and DMEM+FA. During maintenance stage, the media were refreshed every 2-3 days.

**Media preparation**

Differentiated CMs were replated on day 15 of differentiation for *in vitro* hPSC-CMs media-induced maturation protocols. The media used were listed below:

| Base medium | Additional supplements |
|---|---|
| RPMI-1640 | 2% B27 with insulin. |
| DMEM no glucose (ThermoFisher, Cat.#11966025) / 10 mM HEPES | 2 mM L-carnitine,<br>5 mM creatine,<br>5 mM taurine,<br>1 mM nonessential amino acids,<br>1 × insulin-transferrinselenium (ThermoFisher, Cat. #41400045),<br>1 × linoleic-oleic acid (Merck Life Science, Cat. #L9655-5ML). |

**Contractile performance and CardioExcyte 96 recordings**

Differentiated hPSC-CMs were replated on day 15 of differentiation for contractility recording over a 7-day window. We plated 50,000 viable cells per well on Vitronectin XF-coated 96-well CardioExcyte NSP-96 plates (Nanion Technologies GmbH, Cat.#201001). Contractile parameters of beating hiPSC-CMs monolayers were recorded using the CardioExcyte 96 system (Nanion Technologies GmbH) with 1 ms time resolution and 1 kH sampling rate. All experiments were conducted under physiological conditions using an incubation chamber part of the system (37 °C, 5% $CO_2$ and 80% humidity). The CardioExcyte NSP-96 plates use a rigid surface with two gold electrodes in each well to measure contractility via continuous impedance measurements on CMs exposed to various culture conditions. The parameter used in this study is the contractile amplitude.

**Immunofluorescence of hiPSC-CM and sarcomere length analysis**

Differentiated hPSC-CMs were replated on day 15 of differentiation. hPSC-CM were seeded at 40,000 cells per 35 mm MatTek glass-bottom dish (P35G-1.0-14-C) and cultured overnight at a 37°C, 5% $CO_2$ incubator. Post-replating, cells were maintained in RPMI+B27 for 48 hours before refreshing with two backbone media RPMI+B27 vs DMEM+FA. Cells were fixed with 4% paraformaldehyde for 15 min at room temperature, washed in PBS, permeabilised with 0.1% Tween-20 for 10 min, and blocked in 10% donkey serum with 0.1% Triton X-100 for 1 h at room temperature. Primary antibody against α-actinin (1:500, mouse monoclonal, Sigma A7811) was incubated overnight at 4°C in blocking buffer, followed by incubation with Alexa Fluor 488-conjugated anti-mouse secondary antibody (1:200, ThermoFisher) for 1 h at room temperature in the dark. Nuclei were counterstained with DAPI (1 µg/mL), and samples were imaged using a Zeiss LSM 880 confocal microscope. Secondary-only controls were included.

Captured images were analysed using ImageJ (https://imagej.net/ij/). Images were converted to binary and processed using the "Analyze Particles" tool to calculate Feret's diameter, providing size distribution of α-actinin+ aggregates. Sarcomere length and width were quantified in 6-7 biological replicated cells per condition based on α-actinin staining to assess their organisation.

**Processing bulk RNA-seq data for gene quantification**

hPSC-CMs cultured in two backbone media over a 7-day window were harvested for RNA preparation and genome wide RNA-seq. RNA-seq samples with 3 biological replicates for each group were aligned to hg38 using STAR aligner[6] with the default option, yielding more than 294M uniquely mapped reads in total with an average of 92.84% mapping rate. Gene-level read counts were quantified using htseq-count[7] using the hg38 RefSeq gene annotation.

**Identifying differentially expressed genes**

Genes with count per million (CPM) of at least 1 in at least 2 samples were kept for further analysis. Read counts were normalised using 'calcNormFactors' function available in edgeR package, with TMM method[8]. Negative binomial generalised linear models were fit. Differentially expressed (DE) genes were identified using false discovery rate < 0.001.

**RNA-seq analysis of myofilament isoforms**

Raw and transcripts per million (TPM), matrices were downloaded from the Gene Expression Omnibus (GEO) database for studies investigating cardiomyocyte maturation using *in vitro* models: GSE113871, GSE114976, GSE115031, GSE115621, GSE124057, GSE129058, GSE137081, GSE152265, GSE152589, GSE187308, and GSE201437. Additionally, adult human heart data (Hahn et al., 2021) were obtained from the Zenodo Data Repository (DOI: 10.5281/zenodo.4114617) for benchmarking. For each of these studies, mean isoform fractions and their standard errors were calculated across control or healthy condition replicates. Sarcomeric protein isoform fractions were calculated as: MYL2 (MYL2 / (MYL2 + MYL7)) and TNNI3 (TNNI3 / (TNNI3 + TNNI1)). While TPM-normalised counts were used for MYL, raw or CPM-normalised counts, depending on availability, were used for TNN to avoid technical bias from TPM length-normalisation due to the large non-translated exon in TNNI1.

### QUANTIFICATION AND STATISTICAL ANALYSIS

CMs on day 15 were used from 3 independent hPSC differentiation replication experiments. Each experiment group was performed with a minimum of three biological replicates. Statistical analysis was performed using GraphPad Prism (v9.3.1) software. To compare two normally distributed groups, a student's *t*-test was performed. For three or more groups and the assessment of one parameter, a One-Way ANOVA statistical test was used. With multiple parameters and three or more groups, a Two-Way ANOVA was used. Significance was determined as $^{*}p < 0.05$, $^{**}p < 0.01$, $^{***}p < 0.001$ and $^{****}p < 0.0001$. Data are presented as mean ± SEM.

### REFERENCES


1. Ewoldt, J., DePalma, S., Jewett, M., Karakan, M.Ç., Lin, Y.-M., Mir Hashemian, P., Gao, X., Lou, L., McLellan, M., Tabares, J., et al. (2024). Induced pluripotent stem cell-derived cardiomyocyte in vitro models: tissue fabrication protocols, assessment methods, and quantitative maturation metrics for benchmarking progress. Version 8 (Dryad). https://doi.org/10.5061/DRYAD.KSN02V7BH https://doi.org/10.5061/DRYAD.KSN02V7BH.

2. Seltmann, S., Lekschas, F., Müller, R., Stachelscheid, H., Bittner, M.-S., Zhang, W., Kidane, L., Seriola, A., Veiga, A., Stacey, G., et al. (2016). hPSCreg—the human pluripotent stem cell registry. Nucleic Acids Research *44*, D757–D763. https://doi.org/10.1093/nar/gkv963.

3. Funakoshi, S., Fernandes, I., Mastikhina, O., Wilkinson, D., Tran, T., Dhahri, W., Mazine, A., Yang, D., Burnett, B., Lee, J., et al. (2021). Generation of mature compact ventricular cardiomyocytes from human pluripotent stem cells. Nat Commun *12*, 3155. https://doi.org/10.1038/s41467-021-23329-z.

4. Pedregosa, F., Varoquaux, G., Gramfort, A., Michel, V., Thirion, B., Grisel, O., Blondel, M., Prettenhofer, P., Weiss, R., Dubourg, V., et al. (2011). Scikit-learn: Machine Learning in Python. Journal of Machine Learning Research *12*, 2825–2830.

5. Xia, X. (2012). Position Weight Matrix, Gibbs Sampler, and the Associated Significance Tests in Motif Characterization and Prediction. Scientifica (Cairo) *2012*, 917540. https://doi.org/10.6064/2012/917540.

6. Dobin, A., Davis, C.A., Schlesinger, F., Drenkow, J., Zaleski, C., Jha, S., Batut, P., Chaisson, M., and Gingeras, T.R. (2013). STAR: ultrafast universal RNA-seq aligner. Bioinformatics *29*, 15–21. https://doi.org/10.1093/bioinformatics/bts635.

7. Anders, S., Pyl, P.T., and Huber, W. (2015). HTSeq—a Python framework to work with high-throughput sequencing data. Bioinformatics *31*, 166–169. https://doi.org/10.1093/bioinformatics/btu638.

8. Robinson, M.D., McCarthy, D.J., and Smyth, G.K. (2010). edgeR: a Bioconductor package for differential expression analysis of digital gene expression data. Bioinformatics *26*, 139–140. https://doi.org/10.1093/bioinformatics/btp616.
